# Phototherapy with physically and molecularly produced light for Alzheimer’s disease

**DOI:** 10.1101/2022.02.10.478818

**Authors:** Shi Kuang, Biyue Zhu, Jing Zhang, Fan Yang, Bo Wu, Weihua Ding, Shiqian Shen, Can Zhang, Chongzhao Ran

## Abstract

Over the past decades, classical drug development approaches for Alzheimer’s disease have yielded limited success, and this futileness has prompted scientists to seek non-classical approaches. In this report, we demonstrated that, with irradiation of LED light or with molecularly generated light (dubbed as “molecular light”) from chemiluminescence probe ADLumin-4, photolabile curcumin analogue CRANAD-147 could change properties, structures (sequences) and neurotoxicity of amyloid beta (Aβ) species in vitro. We further demonstrated that, with the assistance from molecular chemiluminescence imaging, the combination of CRANAD-147/LED or CRANAD-147/ADLumin-4 (molecular light) could slow down the accumulation of Aβs in transgenic 5xFAD mice in vivo. Due to the unlimited capacity of tissue penetration of molecular light in vivo, phototherapy with the combination of photolabile Aβ ligand and molecular light has great potential as an alternative approach for AD drug discovery.

## Introduction

Photomedicine is a branch of medicine that utilizes light for treatment and diagnosis and it has been widely applied in numerous medical fields ^[1]^. The light for photomedicine is always physically generated from various devices, such as incandescent light bulbs, laser and light emitting diodes (LED). However, light can also be generated molecularly from chemiluminescent, bioluminescent compounds and Cerenkov luminescent tracers, such as luciferins, luminol, and ^18^F-FDG ^[2]^. Here, we dubbed the molecularly generated light as “molecular light”. “Molecular light” has been widely used for imaging purposes, particularly for preclinical animal studies ^[2a, 3]^. “Molecular light” has also been considered as “self-illuminated” molecules ^[4]^. Compared to physically generated light, molecular light has rarely been used for the purpose of phototherapy. We demonstrated that molecular light from Cerenkov luminescence of ^18^F-FDG could be used for in vivo uncaging reaction ^[2b]^. Recently several reports suggested that molecular light can sensitize photosensitizers to generate ROS in photodynamic therapy ^[2a]^.

For phototherapy, physically produced light can only be shone externally and has limited capacity for tissue penetration, and it can’t be delivered to target of interests beyond shallow locations, such as skin and surfaces of internal cavities ^[5]^. By contrast, “molecular light”, which behaves like molecules, can be delivered to targets that can specifically bind to the molecules; therefore, molecular light has nearly limitless tissue penetration. For phototherapy with physically generated light, we can externally shine a large quantity of photon flux, but only few of the photons can reach the target-of-interest inside the body ^[5]^. By contrast, the total photon flux from molecular light is considerably smaller, but it can be delivered to the target-of-interest as close as to a few nanometers to generate chemi-or bio-luminescence resonance energy transfer (CRET or BRET) if the energy from the molecular light matches with the absorbance of the target-of-interest. The distance between target-of-interest and molecular light *vs* physically product light (nanometer *vs* centimeter) is approximately ∽10^6^-folds closer (Fig.1A). In these regards, we speculate that molecular light may have equal or better capacity to initiate photoreactions in vivo.

**Fig.1.**
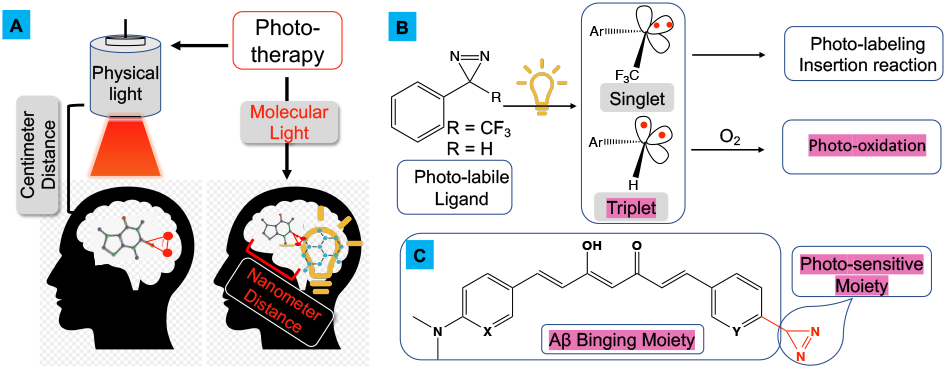
Illustration of phototherapy with physically produced light and molecularly generated light (“molecular light”). Molecular light can be delivered as molecules to avoid tissue penetration limitations of light; B) Reaction mechanism and photoreaction of diazirines; and C) Design of photo-labile curcumin analogue CRANAD-147. D) Synthetic route for CRANAD-147.

Physically produced light has been widely used for photodynamic therapy (PDT) and for initiating photochemistry of photolabile compounds; however, the application of “molecular light” for such purposes has been rarely reported ^[2a]^. In this work, as a proof-of-concept study, we demonstrated that, in the presence of photolabile curcumin, “molecular light” from chemiluminescent compounds could attenuate neurotoxicity of amyloid beta (Aβ) in vitro and in vivo.

Aβ peptides are generated from amyloid peptide precursor (APP) by cleavage with β-and γ-secretases ^[6]^. In the brains of Alzheimer’s disease (AD) patients, these soluble peptides are gradually accumulated and consequentially converted into alternative tertiary conformations, which lead to self-assembly to generate amyloid protofilaments and later ultimately assemble into amyloid plaques ^[7]^. Currently, most AD drugs under preclinical and clinical studies are related to Aβ targeting, such as reducing Aβ productions, preventing Aβ aggregation and promoting Aβ clearance ^[8]^. However, strategies that can reduce Aβ neurotoxicity via altering the structures of amino acids in Aβ sequence has been rarely reported. In this report, we demonstrated that, under irradiation of LED light, photolabile curcumin analogue CRANAD-147 could lead to structural changes of amino acids of Aβ peptides and consequential attenuation of Aβ’s neurotoxicity. Moreover, we showed that “molecular light” from chemiluminescence compound ADLumin-4 could be used to replace the LED light for altering the structures of amino acids. Lastly, we demonstrated that, under LED irradiation, CRANAD-147 could reduce Aβ burdens in transgenic 5xFAD mice after one month of treatment. It is known that light from external irradiation has a limited capacity of tissue penetration for in vivo studies. To overcome this problem, we hypothesized that chemiluminescent ADLumin-4 could be used as a deliverable molecular light source to replace external LED irradiation for in vivo therapeutic treatment. Indeed, we found that the combination of CRANAD-147 and ADLumin-4 could significantly reduce Aβ burdens in 5xFAD mice. In addition, we demonstrated that molecular imaging with our previously reported chemiluminescence probe ADLumin-1 could be used to monitor the efficiency of phototherapy under LED light and ADLumin-4 treatment ^[3a]^.

## Results

### 1. Design and synthesis of CRANAD-147

Abundant evidence suggested that curcumin could have a high binding affinity to Aβs ^[9]^, and we have reported a series of curcumin-base imaging probes CRANAD-Xs for various Aβ species, including near-infrared fluorescence (NIRF) probes ^[10]^, two-photon probes ^[11]^, as well as ROS probes ^[12]^. In our previous studies, we showed evidence that curcumin analogues CRANAD-3, -17-, and -58 could bind to Aβ 13-20 (HHQKLVFF) fragment ^[10]^. Similarly, Masuda et al also demonstrated that curcumin itself could interact with this Aβ fragment ^[13]^. Based on these results, we propose to use the curcumin scaffold as an anchor to target Aβ species.

Over the past decades, numerous photolabile reactions have been broadly explored in photochemistry and biological studies^[14]^, and several photolabile moieties have been widely applied ^[14-15]^. Among these photolabile moieties, diazirine is very attractive due to its excellent light sensitivity and its small size as a tag ^[16]^. The first diazirine photoreaction was reported very early in 1973 ^[16-17]^, and this photosensitive reaction has been applied in numerous biological studies ^[16]^. In recent years, diazirine has emerged as an attractive photo-chemistry tool for various purposes, particularly for photo-crosslinking, bioconjugations, photo-labeling, identifying target engagement (fishing binding targets for drugs) and protein-protein interactions ^[16]^. Previously, the diazirine moiety was installed at the non-terminal position of an aliphatic chain. In this case, to activate this group, short-wavelength light is needed. In addition, the photo-reactivity of the aliphatic diazirine is low and a long-time irradiation is necessary ^[18]^. The short-wavelength and low sensitivity are troublesome for biocompatibility. Considering the above facts, we propose to introduce the diazirine moiety into the long π-π conjugated curcumin system to extend its wavelength of photoreaction and increase its photosensitivity (long π-π conjugation lowers its required energy for activation). In principle, diazirine derivatives have potential as photosensitizers via the triplet state of carbine ^[16]^ (Fig.1B). However, this feasibility has been rarely explored. In these regards, we designed CRANAD-147 via attaching the small diazirine moiety to the terminal of the curcumin scaffold (Fig.1C).

Recently, Reboul’s group reported a series of terminal diazirines that could be used for para-hydrogen induced hyperpolarization ^[19]^. In their synthesis study, they found the phenyliodine (III) diacetate (PIDA) could catalyze imine oxidation to generate the diazirine in the presence of NH_3_. We explored this method and successfully synthesized CRANAD-147 via a phenyl-imine intermediate (Fig.1D). To the best of our knowledge, CRANAD-147 is the first diazirine at the terminal of a long conjugated aromatic system.

### 2. Photo-sensitivity of CRANAD-147

We found that CRANAD-147 could be highly efficiently activated under the 470 nm visible light and the half lifetime t_1/2_= 4.6 min. upon irradiation with a 1-min. duration. HPLC analysis showed a new peak with higher polarity (SI Fig 1). Mass spectrometry results indicated that m/z=377.2 was the major peak, which is likely originated from methanol (solvent) attacking the carbene intermediate. Another apparent peak m/z = 388.2 could be ascribed to the photoaffinity reaction with solvent acetonitrile (SI Fig.2). A product of water-carbene reaction also could be found (m/z=363.2). The multiple products from the photo-reaction is consistent with the nature of high reactivity of carbene intermediates.

### 3. CRANAD-147’s binding with Aβ aggregates and Aβ property changes after light irradiation

We first studied CRANAD-147’s binding affinity toward Aβs. Upon binding to Aβ aggregates, it displayed a modest fluorescence increase (11.3-fold) after 60 minutes incubation. Meanwhile, the quantum yield was increased from 0.003 to 0.026. The time-course of fluorescence increase upon Aβ binding showed that a binding equilibrium was reached within 60 minutes (Fig.2A). We also performed fluorescence titration with Aβ aggregates and obtained a good binding with *K*_*d*_ value of 2.89 ± 0.43 μM (Fig. 2B), which is similar to the binding of curcumin itself with Aβ aggregates ^[20]^. These results suggested that the introduction of diazirine moiety was compatible for curcumin scaffold’s binding to Aβs.

**Fig.2.**
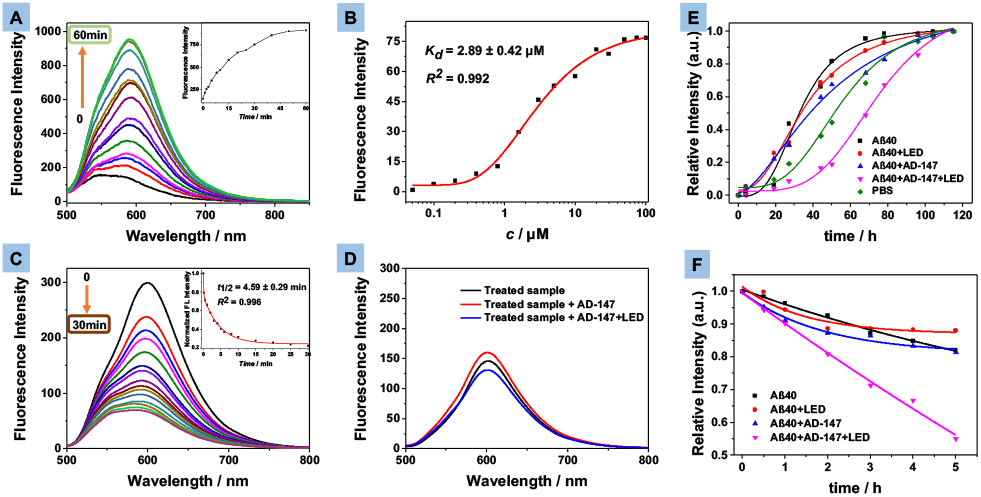
A) Fluorescence spectra of CRANAD-147 after incubation with Aβ aggregates for different time durations. B) Binding constant Kd measurement of CRANAD-147 with Aβ aggregates. C) Fluorescence intensity and time-course of CRANAD-147 with Aβs after LED irradiation. D) Fluorescence spectra of CRANAD-147 after LED irradiation and after adding fresh CRANAD-147 for rescuing study. E) Time-course of seeding with Aβ aggregates from different treatments. F) Time-course of proteinase K degradation of Aβs (ThioT fluorescence). The degradation can be accelerated by photoreaction of CRANAD-147 after LED irradiation.

We reasoned that the diazirine group would be activated under light irradiation and produce highly reactive carbene intermediates that can react with the nearby Aβ peptides that are binding with the curcumin scaffold. To verify this speculation, we first recorded the fluorescence intensity change of CRANAD-147 in the presence of Aβ aggregates over time (Fig.2C) after exposing the mixture under a LED array light (4 W/cm^2^, 470 nm) for 2 seconds for each time point. We observed significant decreases of intensity upon light irradiation with a half lifetime of 4.59 minutes (for 2.5 μM CRANAD-147 with 500 nM Aβ aggregates). We also performed control experiments with thioflavin T (ThioT), which can bind Aβ aggregates but not photolabile. Although we also observed fluorescence intensity decreases, its half-lifetimes (13.8-minutes) were much longer than that of CRANAD-147 (SI Fig.3). These results suggested that CRANAD-147 was photolabile and was converted to new products.

To further investigate the reaction between CRANAD-147 and Aβs, we carried out fluorescence rescue experiments via adding fresh CRANAD-147 into the photo-reaction solution. Interestingly, we only observed less than 10% intensity recovery, suggesting that, during the irradiation, the properties of Aβ aggregates were irreversible altered (Fig.2D). In this regard, we investigated whether the photo-reaction can lead to property changes of Aβ aggregates. We conducted seeding experiments with CRANAD-147/Aβ/light and CRANAD-147/Aβ/dark, and found that the seeds from the light-irradiation group had much poorer capacity for inducing Aβ aggregation (half-time of aggregation: 33.6 vs 68.2 hours) (Fig.2E). In addition, we conducted experiments with Proteinase K digesting to examine whether the photo-reaction could have effects on degradation of Aβ aggregates, and we found that the irradiated Aβ/CRANAD-47 mixture showed much faster degradation (Fig.2F). Taken together, the above results suggested that the photoreaction could altered the properties of Aβ aggregates.

It was also reported that the catechol could increase photon-induced free radical reaction efficiency and decrease oxidative damage ^[21]^. In this regard, we added catechol as a cocatalyst and performed the same photoaffinity reaction. As we can see in SI Fig.4, after light irradiation, the fluorescence was still vanishing, suggesting that adding catechol had no apparent effect in our system because CRANAD-147 is strongly binding into the Aβ hydrophobic pocket that insulates catechol for entering the reaction system. This result also indicated that the photoreaction of Aβ peptide was only confined inside the Aβs, and would not impact the nearby proteins.

We further investigated the property changes of Aβs using SDS-PAGE gel electrophoresis. After running the gel, we imaged the gel on an IVIS imaging system after silver staining (Fig.3A,B) and western blotting (SI Fig.5). From the images, several new bonds that migrated slowing than the control group could be unambiguously identified, suggesting that Aβs were crosslinked while being irradiated in the presence of CRANAD-147. According to the standard gel ladder, these bands could be ascribed to soluble monomers, dimers, trimers, and hexamers. In addition, time-dependent irradiation also indicated that, with the light dose increase, the intensity of bands of the soluble oligomers were remarkably increased (Fig.3C, SI Fig.6).

**Fig.3.**
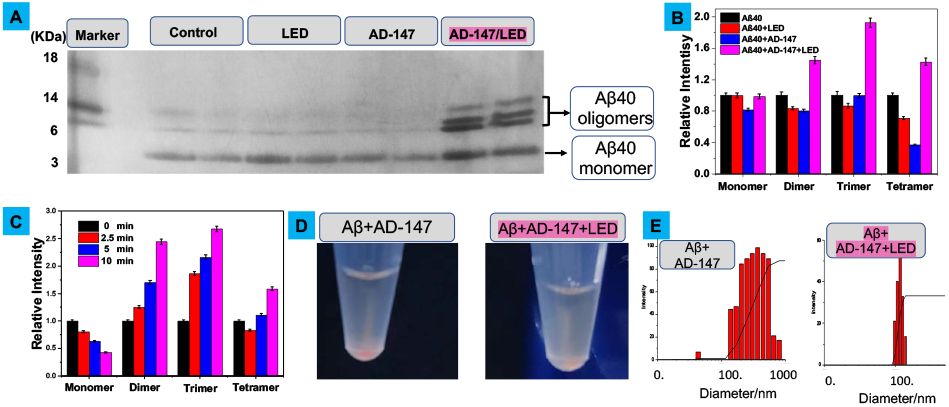
A) SDS-Page gel of Aβs after different treatments. B) Quantitative analysis the bands in (A). C) Quantitative data of monomeric and oligomeric Aβs after different irradiation time with CRANAD-147. D) Photographs of mixtures of Aβs and CRANAD-147 with/without LED irradiation (left: more precipitates). E) Particle sizes of the mixtures.

Interestingly, we observed that there was less precipitation from the tube under irradiation with CRANAD-147, compared to the tube without irradiation (Fig.3D), suggesting the irradiation resulted in better solubility of Aβs, likely due to polarity increases that caused by oxidation of amino acid residues. Consistent with this phenomenon, we found that the particle sizes were much smaller from the irradiation samples (Fig.3E). This modification could be the reason why the irradiated Aβs can be easier degraded by proteinase K.

### 4. Identified structural change of amino acids of Aβ Peptides

We first characterized the Aβ changes with LC-MS. We found several clusters of MS peaks. As shown in Fig.4A, cluster of m/z = 860.8, 870.0, 872.9 and 876.8 represents Aβ40 monomer (5+ charges), oxidized Aβ40 monomer with one, two and three oxygens (5+ charges), respectively. Similarly, clusters of m/z= 1083.2, 1087.2 and 1091.2, m/z = 1443.9, 1449.4 and 1454.6 represent Aβ monomers and oxidized monomers with four (4+) and three charges (3+) respectively. All the results suggested that the monomeric bands in Fig.3A contained plentiful of oxidized monomers. Interestingly, we also found clusters of oligomerized Aβs and oxidized oligomers (Fig.4B, tetramers and octamers). We further used MALDI-MS to confirm the existence of irreversible oligomerized Aβs, evident by peaks of dimer (m/z=8653.7), trimer (m/z=12990.9), and larger oligomeric Aβs. Interestingly, we noticed that m/z=17348.7, 26030.5, 30361.1, 34693.9 were not exact masses of the corresponding oligomeric native Aβs (Fig.4C). Instead, for each Aβ unit, an 8-Dalton higher mass could be found when it is compared to the native Aβ40 peptide, indicating half of the Aβ units were oxidized (+16 Dalton) in these oligomers (Fig.4C). As expect, we could not find such characteristic oxidized peaks and oligomerized peaks from the non-irradiated control groups (SI Fig. 7, 8, 9).

**Fig.4.**
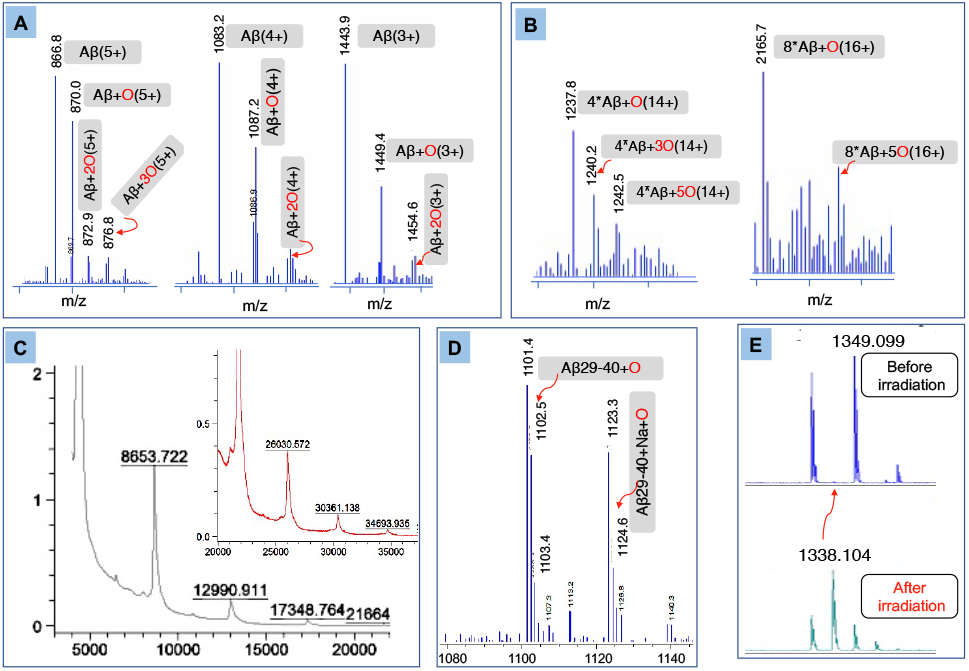
A) Mass spectra (MS) of oxidized monomeric Aβs with different charges. B) MS of oxidized oligomeric Aβs. C) MALDI-TOF MS of oligomeric Aβs and their oxidized products. D) MS of oxidized Aβ29-40. E) MALDI-MS of oxidized Aβ6-16 fragment.

To further investigate the cross-linked products, the sample mixtures were digested by the trypsin and analyzed by LC-MS. It is well-documented that the trypsin will selectively cleavage C-terminal of Arginine (R) and lysine (K) ^[22]^. In the case of Aβ40 peptide, the possible trypsinized fragments include Aβ1-5 (DAEFR), Aβ 6-16 (HDSGYEVHHQK), Aβ17-28 (LVFFAEDVGSNK), and Aβ29-40 (GAIIGLMVGVV). From LC-MS of the trypsinized mixture, as shown in SI Fig.10, fragments Aβ1-5 and Aβ17-28 could be easily found since their m/z signatures are apparently identifiable. M/z =637.2 and 319.2 (2+), and m/z =1325.4 and 663.3 (2+) were corresponding to Aβ1-5 and Aβ17-28 respectively. Aβ29-40 also could be found (m/z=1085.6 and 543.3) (Fig.4D); however, interestingly, several high molecular weight peaks could also be observed. After carefully analysis, we assigned the peaks to Aβ29-40+O (m/z =1102.5, m/z =1123.4 plus sodium ion) (Fig.4D), suggesting that Aβ29-40 fragment was oxidized and this oxidation is in accordance with the above MALDI-MS results. Methione-35 is the likely amino acid to be oxidized since it is the most vulnerable residue for oxidation ^[23]^, which has been reported that oxidation of Met-35 to the sulfoxide lead to less neurotoxicity of Aβs by altering production of toxic Aβ oligomers ^[23c, 24]^. Nonetheless, fragment Aβ6-16 was difficult to be identified after irradiation, suggesting some structural changes occurred for this fragment. Likely the Histidine-13 or -14 was oxidized and degraded ^[23, 25]^.

To further find evidence, we used high resolution MALDI-MS to investigate the trypsinized mixture without liquid-chromatography separation. Consistent with the data from LC-MS, as shown in SI Fig.11, the trypsinized fragments Aβ1-5, Aβ17-28, Aβ29-40 and Aβ29-40+O could be found. Notably, there was one new peak (m/z =1338.1) could be easily observed for the irradiated sample. Given that histidine could be photo-oxidized into Aspartic acid (D) or Asparagine (N) and tyrosine could also be photo-oxidized ^80,81^, we assigned m/z= 1338.1 to modified Aβ6-16 fragment NDSGY(+O)EVNH(+2O)QK, in which two histidines (H6 and/or H13 or H14) were converted into Asn and one histidine and tyrosine were oxidized. Taken together, both LC-MS and MALDI-MS data provide evidence that the structure of Aβ6-16 was changed. Collectively, our MS results suggested several structural modifications (covalent cross-linking/oligomerization, oxidation of amino acid residue) of Aβ peptides under LED irradiation with CRANAD-147. These structural modifications are similar to previously reported results from Kanai group ^[26]^. However, they didn’t identify which amino acids or fragments were modified.

## 5. Replacement of LED light with molecular light for in vitro studies

In this report, we propose to use the molecularly generated light from chemiluminescence probes to replace the LED light. Our previous report showed that ADLumin-1 is an excellent Aβ targeting chemiluminescence probe ^[3a]^. It can transfer the energy from itself to the near infrared fluorescence (NIRF) probe CRANAD-3 via chemiluminescence resonance energy transfer (CRET). Herein, given that the emission between ADLumin-1 and CRANAD-147 only have a 10-nm separation, suggesting that ADLumin-1 and CRANAD-147 is not an ideal CRET pair. In order to solve this problem, we synthesized ADLumin-4 (Fig.5A), which has a shorter wavelength of emission. Compare with the ADLumin-1, the ADLumin-4 has only one double bond and its emission is 40 nm shorter, and this emission has much better a spectral overlap match with the absorption spectrum of CRANAD-147. Indeed, the energy transfer efficiency was quite high between ADLumin-4 and CRANAD-147, evident by the low signal at 450nm after CRETing (Fig.5B, red line). We further investigated the CRET induced photoreaction of CRANAD-147 in the presence of Aβs. To this end, CRANAD-147 (1 μM) was first incubated with Aβ aggregates for 60 min. After that, different concentrations of ADLumin-4 were added. Next, we used the SDS-PAGE gel electrophoresis to analyze the products. As shown in Fig.5C,D, the oligomeric bands could be easily observed from the CRANAD-147/ADLumin-4 pair, which is consistent with the results from LED irradiation of Aβ/CRANAD-147. These results suggested that molecular light from ADLumin-4 could be used to replace LED light for in vitro studies. We also performed proteinase K hydrolysis with Aβ/CRANAD-147/ADLumin-4, and found that, similar to LED irradiation, ADLumin-4 could accelerate the hydrolysis of the aggregated Aβs (Fig.5E). Lastly, we performed LC-MS analysis for the mixture of CRANAD-147/ADLumin-4/Aβs, and found oxidized monomers (m/z=1449.6), tetramers (m/z=1237.9), octamers (m/z=2165.1) (Fig.5F), which were similar to the results from CRANAD-147/LED/Aβs mixture (Fig.4A,B). Taken together, our data (SDS-Page gel, proteinase K and LC-MS) suggested that molecular light from ADLumin-4 could initiate photo-oxidation reaction in vitro in the presence of Aβs.

**Fig.5.**
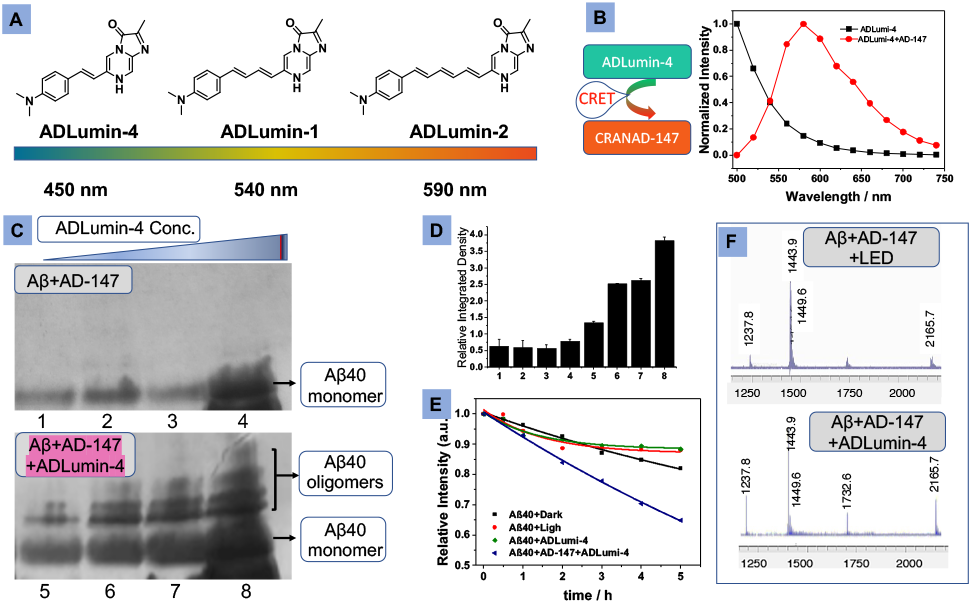
A) Structures and maximum emissions of ADLumin-Zs. B) CRET between ADLumin-4 and CRANAD-147 suggested high transfer efficiency. C, D) SDS-Page and quantitative analysis of Aβ bands (upper: controls of without LED) with increased concentrations of ADLumin-4. E) Time-course of proteinase K degradation of Aβs (ThioT fluorescence). F) MS of oxidized monomeric and oligomeric Aβs with LED (upper) and with molecular light from ADLumin-4 (bottom).

## 6. CRANAD-147 attenuated Aβ’s cell toxicity with LED irradiation and molecular light

We evaluated the relationship between Aβ property changes and its toxicity. First, we examined cell viability with different treatments of Aβ oligomers using an adenosine triphosphate (ATP) assay kit. ATP can be used to measure cell proliferations and cell cycle dynamics, and its concentration decreases if cell viability decreases ^[27]^. As shown in Fig.6A, we found that the viability result from SHY-5Y cells, a neuroblastoma cell line, were apparently increased from the Aβ/CRANAD-147/LED group, compared to the group only with Aβ or Aβ/CRANAD-147. Similarly, after incubating with ADLumin-4 and CRANAD-147 together with Aβs, the cell viability exhibited a noticeable increase, compared to other groups. Next, the neurotoxicity of Aβs was evaluated using an adenylate kinase assay method. Adenylate kinase is an essential intracellular enzyme ^[28]^. When a cell is damaged, the kinase leaks through the cell membrane into the medium, Therefore, measurement of the medium concentrations of adenylate kinase can be used to report the cell toxicity. As shown in Fig.6B, the Aβs/dark group showed the highest toxicity towards the neuroblastoma cells, while the Aβs/CRANAD-147/LED group and Aβs/CRANAD-147/ADLumin-4 groups showed decreased toxicity. Considering SHY-5Y is a cancer cell line, it may not be a good model to mimic normal neuronal cells. In this regard, we used 3D organoids of induced pluripotent stem cell (iPSc) derived neurons to perform the above ATP cell viability studies. The results showed that CRANAD-147/LED and CRANAD-147/ADLumin-4 groups could significantly reduce Aβ’s toxicity (Fig.6C). We further performed TUNNEL assay with the 3D organoid slices via imaging with ApopTag(®) kits, and the results suggested that CRANAD-147/LED and CRANAD-147/ADLumin-4 groups had less apoptosis (Fig.6D-G).

**Fig.6.**
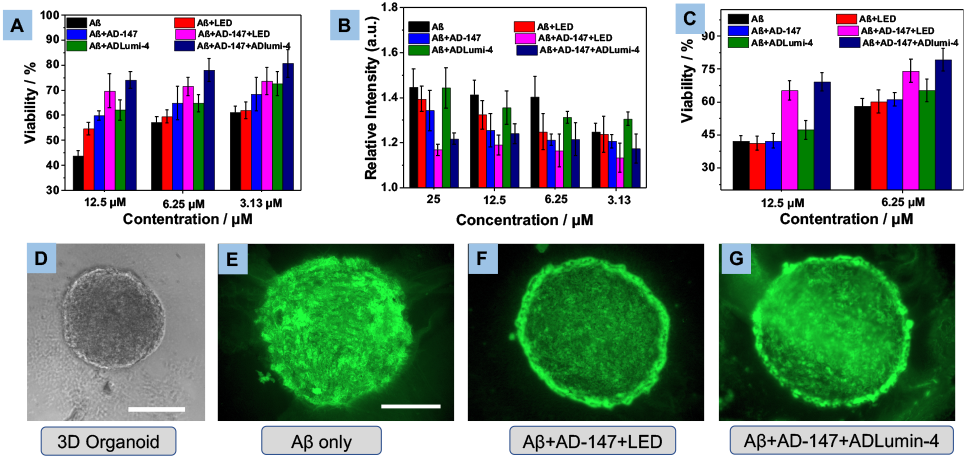
A) Cell viability assays by ATP kit of SHY-5Y cells with different Aβs. B) Aβ neurotoxicity reported via adenylate kinase assays with SHY-5Y cells. C) Cell viability assessment with ATP method for 3D organoids from iPS cell cultures. D-G) Representative images of 3D organoid slices visualized by ApopTag(®) for apoptosis induced by Aβs. Scale bar: 500 μm.

## 7. In vivo therapeutic studies with CRANAD-147

Based on the above promising results in vitro, we investigated whether photolabile CRANAD-147 has therapeutic effects under LED light and with molecular light. Firstly, we performed mimic studies in a biologically relevant environment. To this end, we used brain homogenates from a wild-type mice to examine whether structural modifications of Aβ could be observed. We externally added Aβs to the brain homogenates, and the CRANAD-147/Aβ group was irradiated with 470 nm LED light for 5 min. As shown in (SI Fig.13), after being treated with the CRANAD-147/Light group displayed an apparent band (band 4) that could not found in all the other groups, suggesting that CRANAD-147 can selectively binding to Aβ to induce changes of its properties in biologically relevant conditions. Similarly, the same band could be observed in the CRANAD-147/ADLumin-4/Aβ group.

To validate whether CRANAD-147 and ADLumin-4 can bind to Aβ species in biologically relevant environments, we incubated the compounds with brain slides of a 5xFAD mouse. As expected, both compounds can label the plaques in the brain (Fig.7A), suggesting that both compounds are specifically binding to Aβ deposits.

**Fig.7.**
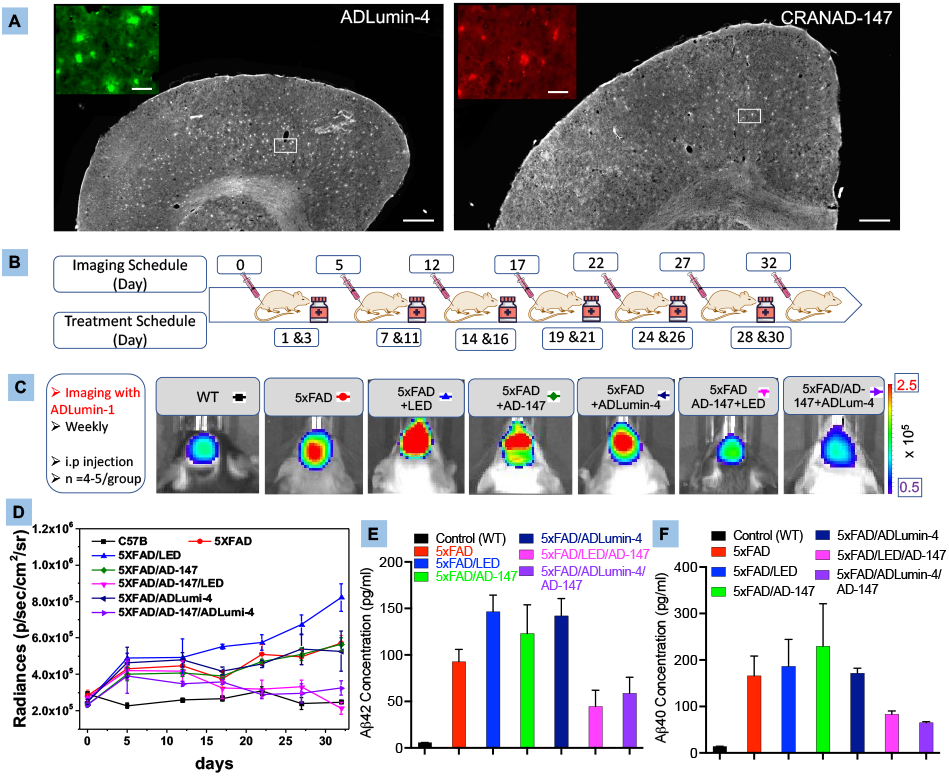
A) Histological staining of brain slides of a 5xFAD mouse (8-month-old) with ADLumin-4 (left) and CRANAD-147(right). Insert: Zoomed-in image of the white box area. B) Diagram of in vivo imaging and treatment schedule. C) Representative images of different groups at Day 32. D) Quantitative analysis of chemiluminescence intensity of images captured at 30 minutes after i.p injection of ADLumin-1. There are significant differences between controls and experimental groups with LED irradiation (pink) or molecular light from ADLumin-4 (purple). E) Concentrations of Aβ42 measured by immunoassays. F) Concentrations of Aβ40 measured by immunoassays.

To investigate whether CRANAD-147/LED and CRANAD-147/ADLumin-4 combination can reduce Aβ burdens in the course ageing, we treated 6 groups of transgenic 5xFAD mice under different treatment as shown in Fig.7B for one month. The mice were administrated CRANAD-147 (2.5 mg/kg) every two days and irradiated with LED light (15 minutes) or treated with ADLumin-4 (10.0 mg/kg) (administrated at the same as CRANAD-147). We monitored the progress every four days with our previously reported chemiluminescence probe ADLumin-1 via *i*.*p*. injection at different treatment time-points. At the meantime, we also used a group of wild type (WT) mice, which do not have overexpressed “humanized” Aβ and thus have no changes of Aβ concentrations, as the control group to make sure our monitoring method is reliable. As shown in Fig.7C,D, during the time-course, there was no significant chemiluminescence intensity increase from the WT group, suggesting that our monitoring method is trustworthy. As expected, we observed a steady increase of intensity from the 5xFAD groups that were not treated with CRANAD-147. Similarly, we observed increasing trends of chemiluminescence signals from groups that were treated with CRAND-147 only (no LED irradiation) or treated with ADLumin-4 only (no CRANAD-147). By contrast, no significant increases of chemiluminescence signals were observed when the 5xFAD mice were treated with CRANAD-147/LED (pink line) or CRANAD-147/ADLumin-4 (purple line), indicating the combinations of CRANAD-147 with light that from LED or ADLumin-4 molecules are necessary to reduce Aβ burdens during treatments.

To validate the above imaging results, we sacrificed the mice at the end of imaging session, and brains were homogenized and extracted with TBST buffers. With the brain extracts, Aβ concentrations were measured via MSD (Meso Scale Discovery) immunoassays ^[29]^. As expected, the groups of CRANAD-147/LED (pink) or CRANAD-147/ADLumin-4 (purple) showed significant lower concentrations of Aβ42 and Aβ40 (Fig.7E,F), which is consistent with the above in vivo imaging data.

## Conclusions

In the report, we presented novel approaches for phototherapy of AD via attenuating Aβ’s neurotoxicity. Likely, the photoreaction with photolabile curcumin changed the properties of Aβ via oxidation the vulnerable amino acids such as methionine and histidines. Remarkably, we demonstrated that molecularly generated light (molecular light) could have similar effects as LED light on the photoreaction of the photolabile curcumin, suggesting that molecular light could have the potential to overcome the limitations of externally applied light. Lastly, we demonstrated that the combination of photolabile curcumin and LED light or molecular light could reduce Aβ burdens during treatment. In summary, we presented new approaches for AD therapy and our studies opened a new avenue for in vivo photochemistry.

## Acknowledgements

This work was supported by NIH R01AG055413, R21AG059134, and R21AG059134 awards (C.R.).

