## Supplemental information for "Phototherapy with physically and molecularly produced light for Alzheimer’s disease"

### Spplemental Information

### Photolabile curcumin with physically and molecularly produced light for attenuating of neurotoxicity of amyloid beta species

Shi Kuang[1], Biyue Zhu[1], Jing Zhang[1], Fan Yang[1], Bo Wu[1], Weihua Ding[2], Shiqian Shen[2], Can Zhang[3], and Chongzhao Ran[1]*

*[1]Athinoula A. Martinos Center for Biomedical Imaging, Department of Radiology, Massachusetts General Hospital/Harvard Medical School, Room 2301, Building 149, Charlestown, Boston, Massachusetts 02129; [2]MGH Center for Translational Pain Research, Department of Anesthesia, Critical Care and Pain Medicine, Massachusetts General Hospital, Harvard Medical School, Boston, MA 02114; [3]Genetics and Aging Research Unit, McCance Center for Brain Health, MassGeneral Institute for Neurodegenerative Disease, Department of Neurology, Massachusetts General Hospital and Harvard Medical School.*

#### Experimental Procedures

##### Materials

All reagents were purchased from Sigma-Aldrich, of reagent grade quality, and were used as received unless stated otherwise. Column chromatography was performed on a glass column slurry-packed with silica gel (60 Å, 40–63 mm; SiliCycle Inc.). Recombinant Aβ peptide (1-40) was purchased from rPeptide. Aβ aggregates for in vitro studies were generated by a slow stirring of Aβ40 in PBS buffer for 3 days at room temperature. $^1$H and $^{13}$C NMR spectra were recorded at 500 MHz and 125 MHz on JOEL spectrometers in $CDCl_3$ solution at room temperature. Liquid chromatography-mass spectrometry (LC-MS) was performed using an Agilent 1200 Series apparatus with an LC/MSD trap and Daly conversion dynode detector with UV detection at 254 nm. Fluorescence measurements were carried out using an F-7100 fluorescence spectrophotometer (Hitachi). The IVIS Spectrum imaging system (PerkinElmer) was used for in vitro imaging.

##### Synthesis and characterization

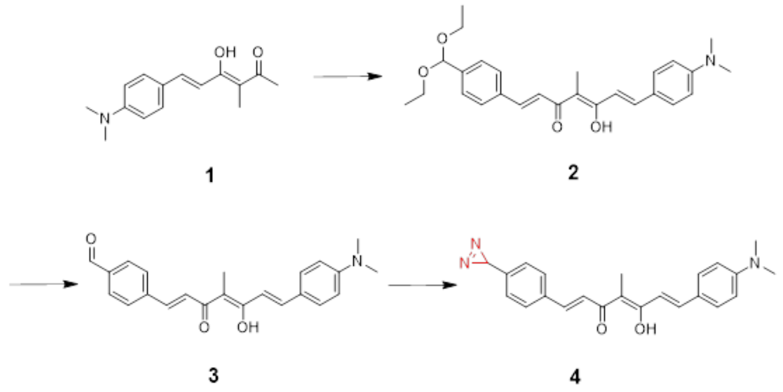


**Scheme S1.** Synthesis of the CRANAD-147.

**Synthesis of 6-(4-(dimethylamino)phenyl)-4-hydroxy-3-methylhexa-3,5-dien-2-one (1)**: $B_2O_3$ (0.7g, 10 mmol) was added into a solution of DMF (8 mL) and the resulting mixture was heated to 120 °C for 30 min until dissolved. To the above mixture, 3-methylpentane-2,4-dione (1.14g, 10 mmol) was added drop-wise. After that, the heating temperature for the mixture was decreased to 110 °C, which was followed by an addition of tributyl borate (4.6 g, 20 mmol). The resulting mixture was stirred for an additional 5 min at 110 °C, followed by an addition of 4-(dimethylamino)benzaldehyde (1.5 g, 10 mmol). The resulting solution was stirred for 5 min, followed by an addition of a solution of 1,2,3,4-tetrahydroquinoline (0.2 mL) and acetic acid (0.4 mL) in DMF (5 mL). The reaction mixture was stirred at 110 °C for 2 h. Upon completion, after cooling to room temperature, the mixture was poured into 400g ice. The resulting solution was filtered and washed with cold water. The precipitate was collected, and the crude product was further purified by flash silica chromatography (HEX: EA=2:1). **Compound 1** was obtained as an orange solid with a 25 % yield. $^1$H NMR (500 MHz, $CDCl_3$): δ 7.60 (d, J = 15.4 Hz, 1H), 7.45 (d, J = 8.9 Hz, 2H), 6.72 (d, J = 15.4 Hz, 1H), 6.67 (d, J = 8.8 Hz, 2H), 3.01 (s, 6H), 2.21 (s, 3H), 1.99 (s, 3H).

**Synthesis of 1-(4-(diethoxymethyl)phenyl)-7-(4-(dimethylamino)phenyl)-5-hydroxy-4-methylhepta-1,4,6-trien-3-one (2)**: Compound 1 (200 mg, 0.8 mmol) was dissolved in 10 mL butanol, followed by an addition of 4-(diethoxymethyl)benzaldehyde (155 μL, 1 eq) and piperidine (120 μL, 1.5 eq). The resulting mixture was refluxed for 24 h. After cooling to room temperature, the solvent was removed under reduced pressure and the residue was washed by water and extracted by ethyl acetate. After removing the solvent under reduced pressure, the crude products were obtained and used for the next step without further purification.

**Synthesis of 4-(7-(4-(dimethylamino)phenyl)-5-hydroxy-4-methyl-3-oxohepta-1,4,6-trien-1-yl)benzaldehyde (3)**: Compound 2 was dissolved with 10 mL dd water and 10 mL $CH_3OH$. The resulting solution was refluxed for 2 h. After cooling to room temperature, the solvent was removed under the deduced pressure and the residue was extracted by ethyl acetate (20 mL*3). The organic phase was collected and then the solvent was removed under reduced pressure. The crude product was purified by flash silica chromatography (HEX: EA=2:1). **Compound 3** was obtained as a red solid with a 20 % yield.

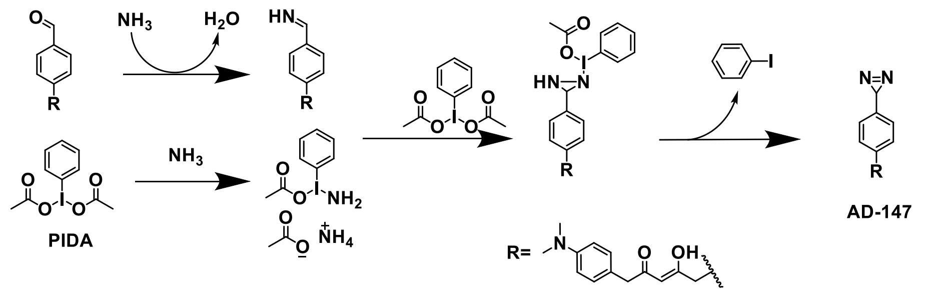


**Scheme S2.** Reaction mechanisum of the terminal diazirine.

**Synthesis of 1-(4-(3H-diazirin-3-yl)phenyl)-7-(4-(dimethylamino)phenyl)-5-hydroxy-4-methylhepta-1,4,6-trien-3-one (4) CRANAD-147**: Compound 3 (23 mg, 0.06 mmol) was added in a round flask that was covered with foil paper, followed by adding of $NH_3 \cdot H_2O$ (14N, 75 μL, 1.05 mmol, 17.5 eq) and $CH_3OH$ (75 μL) at 0 °C. The flask was flushed by Argon gas and the resulting mixture was stirred for 30 min at 0 °C. Then (diacetoxyiodo)benzene (50 mg, 0.15 mmol, 2.5 eq) was added into the flask that was under an inert atmosphere of Argon. The resulting solution was stirred at 0 °C for another 4 h and the reaction mixture was slowly warmed to room temperature overnight. Upon completion, the reaction mixture was concentrated under reduced pressure and the residue was purified by flash silica chromatography (HEX: EA=2:1) under dark. **Compound 4 (CRANAD-147)** was obtained as a red solid with a 28 % yield. 1H NMR (500 MHz, $CDCl_3$): δ 7.67 (d, J = 15.4 Hz), 7.62 (d, J = 15.7 Hz), 7.48 (d, J = 8.7 Hz), 7.43 (d, J = 8.8 Hz), 6.91 (d, J = 15.3 Hz), 6.73 – 6.65 (m), 6.63 (dd, J = 12.1, 3.2 Hz), 4.09 (q, J = 6.9 Hz), 3.04 (s), 3.02 (s), 2.14 (s), 1.44 (d, J = 6.9 Hz).

**Binding affinity Kd measurement and binding kinetics**: For binding affinity Kd: to a solution of CRANAD-147 (1.0 μM) in PBS, different concentrations of Aβ40 aggregates was added, and the fluorescence intensity was recorded until the intensity was not changed. The recorded peak values were fitted by GraphPad with one-site binding model to provide the binding affinity Kd. For time-dependent binding kinetics, CRANAD-147 (1.0 μM) and Aβ40 (500 nm) were mixed in the PBS and the fluorescence emission intensity was recorded every 1 min. for 60 minutes.

**Photoreaction of CRANAD-147:** To a solution of CRANAD-147 (500 nM) in PBS, 250 nM (0.5 eq), 500 nM (1 eq), 1.0 μM (2 eq), 2.5 μM (5 eq) Aβ 40 aggregates was added and incubated for 1 h. The resulting solution was then irradiated by 470 nm LED light (4 $mW/cm^{-2}$, 30 s), and the fluorescence intensity was recorded every 30 s. After that, the data were recorded and fitted by GraphPad software. After the experiments, the samples were kept in the dark overnight. For rescuing experiments, different concentrations of fresh AD-147 were added into the above samples, and the emission intensity was recorded again. To check whether the emission intensity oft he above samples changes after irradiating by the 470 nm again (4 $mW/cm^{-2}$, 30 s), the emission intensity was recorded again after the addition of fresh solutions of CRANAD-147. Fort he photoreaction experiments, Thioflavin T (ThT) was used as a control probe that is not photosensitive.

**Seeding experiments with different Aβ40 aggregates**: The Aβ40 aggregates (300 μL, 25 μM) were treated as following:
Group 1: The Aβ40 aggregates was kept in a dark environment;
Group 2: The Aβ40 aggregates was irradiated by 470 nm LED light (4 mW/cm$^{-2}$, 60 s);
Group 3: The Aβ40 aggregates was mixed with CRANAD-147 (1 eq) and the resulting mixture was kept in a dark environment;
Group 4: The Aβ40 aggregates was mixed with CRANAD-147 (1 eq) and then was incubated for 1 h. Then the resulting mixture was irradiated by 470 nm LED light (4 mW/cm$^{-2}$, 60 s).

After the treatments, all these samples were added to a solution of Aβ40 monomers (6 mL. 25 μM) separately. In every time-point, a 500 μL sample was pipetted out, followed by the addition of ThT (500 nM). Then the emission intensity was recorded at 495 nm (Ex = 435nm). The emission intensities were recorded twice per day until no significant intensity change could be observed. The emission peak intensities were normalized by the last data for each group.

**Protease K digest experiments:** Six groups of Aβ40 aggregates in PBS (3.0 mL, 5 μM) were treated as below respectively.
Group 1: The Aβ40 aggregates was kept in a dark environment;
Group 2: The Aβ40 aggregates was irradiated by 470 nm LED light (4 mW/cm$^{-2}$, 60 s);
Group 3: The Aβ40 aggregates was mixed with CRANAD-147 (1 eq) and the resulting mixture was kept in a dark environment;
Group 4: The Aβ40 aggregates was mixed with CRANAD-147 (1 eq) and then was incubated for 1 h. Then the resulting mixture was irradiated by 470 nm LED light (4 mW/cm$^{-2}$, 60 s);
Group 5: The Aβ40 aggregates was mixed with mixed with ADLumin-4 (10 eq);
Group 6: The Aβ40 aggregates was mixed with CRANAD-147 (1 eq) and then was incubated for 1 h, followed by the addition of ADLumin-4 (10 eq).

After these treatments, all these samples were incubated overnight. Then protease K (0.1 mg/mL) was added to each sample and all of these samples were incubated at 37 ℃° humidity incubator. The Aβ aggregates concentrations were measured by the ThT, and the data was recorded every 30 min.

**SDS-Page gel electrophoresis:** The samples of Aβ40 aggregates were treated as same as the above six groups. The treated samples were loaded into 12-20 % SDS-Page gel with a volume of 20 μL/well. The gel was run with 200 V for 50 min. Then the gel was visulalized by silver stain kit, and the image was captured by the IVIS. For the western blot experiment, the primary antibody was 6E10.

**MALDI-TOF:** The samples of Aβ40 aggregates were treated as same as the above six groups. Then the treated samples were diluted with the HFIP (Hexafluoro isopropanol) (5 μM final concentration) to dissociate no-covalent binding between Aβs.

**Molecular light for CRET between ADLumin-4 and CRANAD-147**: CRANAD-147 (50 μM) and ADLumi-4 (50 μM) was mixed in a DMSO solution, and the chemiluminescent emission spectrum was recorded on an IVIS imaging system with filters of 20nm-band width from 500nm to 700nm. For CRET with Aβs, CRANAD-147 (25 μM) was added to a solution of Aβ40 oligomers (25 μM) in PBS buffer (pH7.4), and the resulting mixture was incubated for 1 h. Then, ADLumi-4 (25 μM) was added to the above mixture and the chemiluminescent emission was recorded by the IVIS imaging system.

**Cell toxicity/viability measurement**: SH-SY5Y cells were seeded in a 96-well plate and incubated for 24 h. To the cell media, different Aβs that were treated as the above groups were added. Then the cells were further incubated 48 h. Upon the completion of culture, ADP kit and ATP kit were used to measure the concentrations of ADP and ATP in the medium separately.

**3D organoids cell culture**: For the 3D brain cell spheroids, about $4\times10^3$ per well SH-SY5Y cells were seeded into the 96-well spheroid bottom plate and incubated in a incubator at 37 ℃° for 3 days. Then the cell were treated as the above six groups respectively. The cell spheroids diameter was recorded every two days. After two weeks of measurement, cell spheroids were dissociated and subjected to ATP assay. For 3D brain organoids, about $10^3$ per well iPSCs were seeded into the 96-well spheroid bottom plate and incubated in 37 ℃° incubators for 3 days. Then the organoids were subjected to treatment of two weeks as the above six groups respectively. After the organoids were fixed and embedded in O.T.C., then the organoids were sliced into slides of 7-micron thickness. The tissue slices were stained with kits of terminal deoxynu-cleotidyl transferase dUTP nick end labeling (TUNEL).

**In vivo therapeutic studies**: 5xFAD mice (12 weeks, female) were randomly separated to 6 groups (n=5 per group). For in vivo imaging, on day 0, 5, 12, 17, 23, 28 and 32, the mice were i.p injected with ADLumi-1 (10 mg/kg, formulation: 2.0 mg/mL, 5% DMSO, 15% Castor oil and 80% saline), and chemiluminescence imaging was performed on the IVIS imaging system with open filter at 30 minutes post the injection, and semi-quabtitive imaging analysis was conducted with average intensity of an ROI of the brain area. For therapeutic studies, the mice were treated on day 1, 3, 6, 8, 10, 13, 15, 18, 20, 22, 24, 26, and 29. After the last imaging session, all the mice were sacrificed and the brains were extracted and frozen in -78°C freezer. For each group, the mice were independently treated as described below:
Group 1: intraperitoneal injection of vehicle of 100 μL of the mixture of 5% DMSO, 15% Castor oil and 80% saline.
Group 2: intraperitoneal injection of vehicle of 100 μL of the mixture of 5% DMSO, 15% Castor oil and 80% saline, 30 min later, the brain areas of the mice were exposed under the 470 nm LED light (2 mW/cm$^2$) for 15 min.
Group 3: intraperitoneal injection of 100 μL of a dose of 2.5 mg/Kg CRANAD-147 (Formulation: 0.5 mg/mL, 5% DMSO, 15% Castor oil and 80% saline).

Group 4: intraperitoneal injection of 100 μL of a dose of 2.5 mg/Kg CRANAD-147 (Formulation: 0.5 mg/mL, 5% DMSO, 15% Castor oil and 80% saline), 30 min later, the mice head were exposed under the 470 nm LED light (2 mW/$cm^2$) for 15 min.
Group 5: intraperitoneal injection of 100 μL of a dose of 10 mg/Kg ADLumi-4 (Formulation: 2.0 mg/mL, 5% DMSO, 15% Castor oil and 80% saline).
Group 6: intraperitoneal injection of 100 μL of a dose of 2.5 mg/Kg CRANAD-147 (Formulation: 0.5 mg/mL, 5% DMSO, 15% Castor oil and 80% physiological saline), 30 min later, the mice were intraperitoneal injected (i.p.) of 100 μL of a dose of 10 mg/Kg ADLumi-4 (Formulation: 2.0 mg/mL, 5% DMSO, 15% Castor oil and 80% saline).

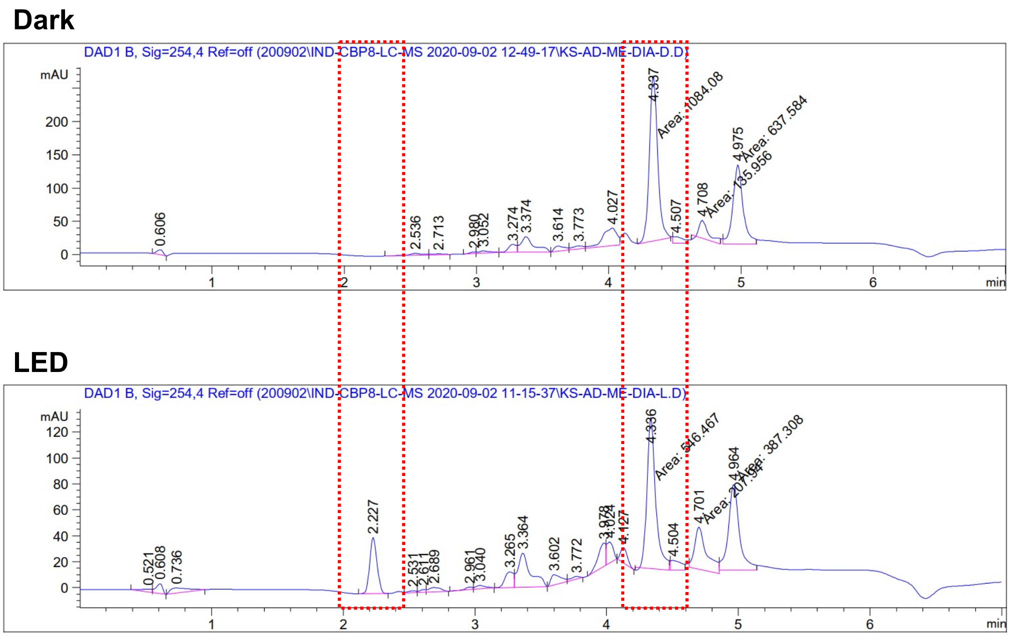


**SI Figure 1.** HPLC of CRANAD-147 before (upper) and after (bottom) LED-irradiation.

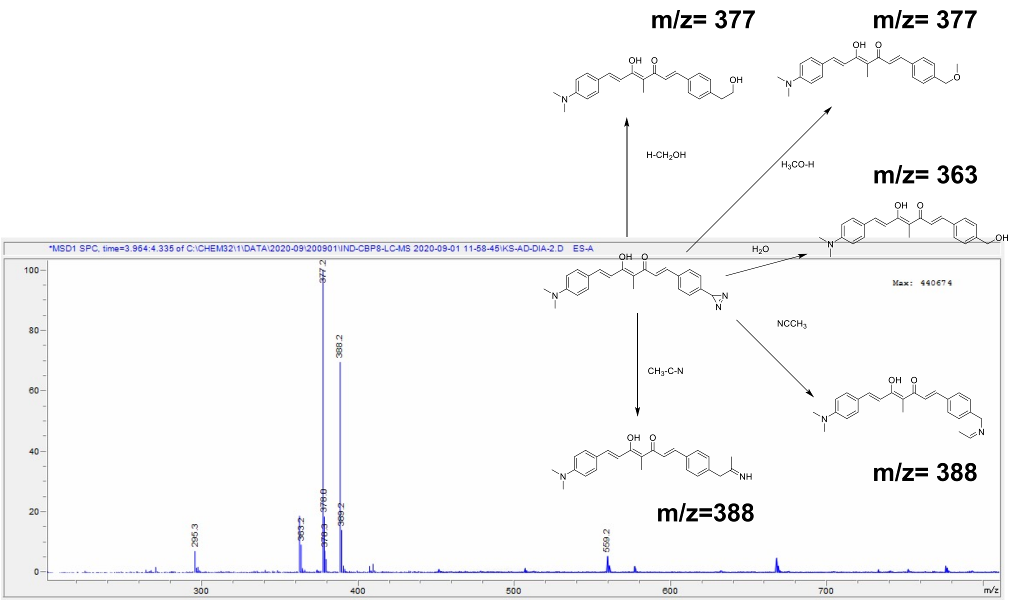


**SI Figure 2.** MS of CRANAD-147 after LED irradiation

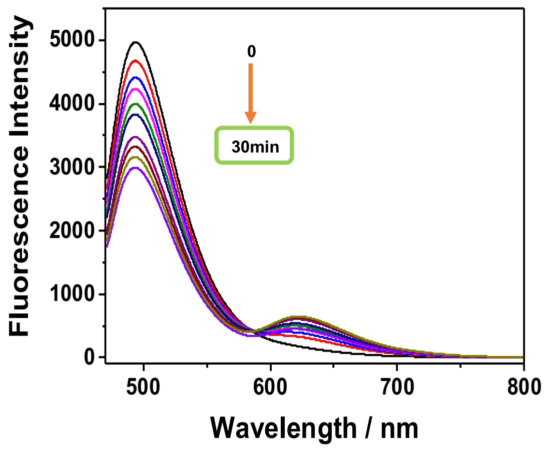


**SI Figure 3.** Fluorescence emission spectra of Thioflavin T after LED irradiation at different time points

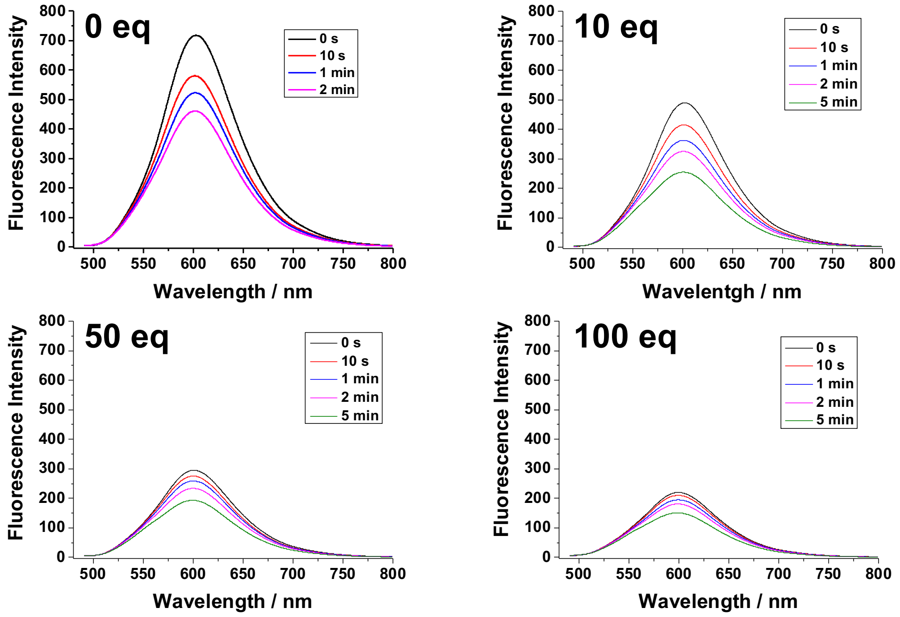


**SI Figure 4.** Fluorescence emission spectra of CRANAD-147 and Aβ40 aggregates with different concentrations of catechol at different time points after LED irradiation.

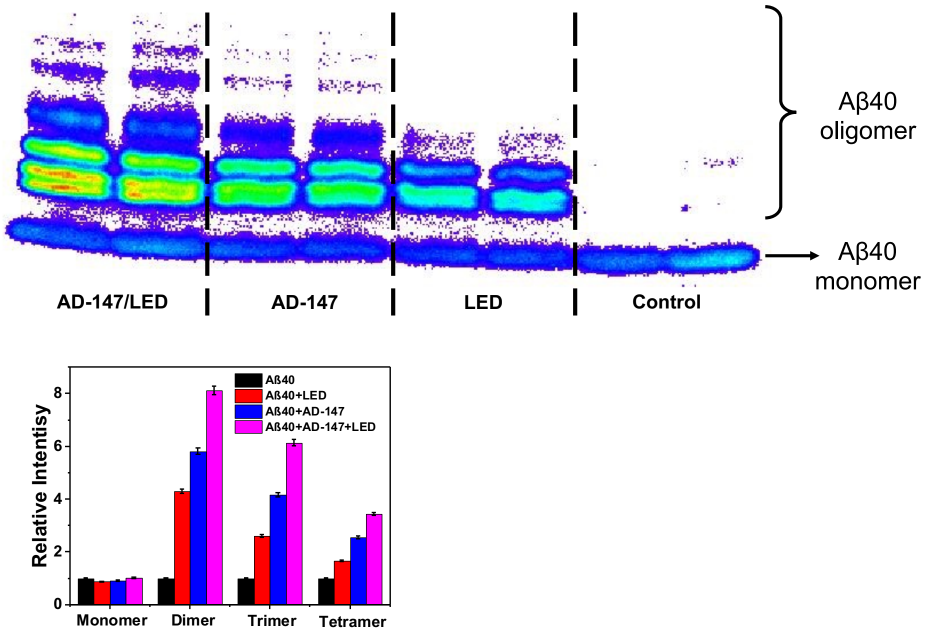


**SI Figure 5.** Western blot of Aβ40 aggregates with different treatments (upper) and quantitative analysis of the bands (bottom).

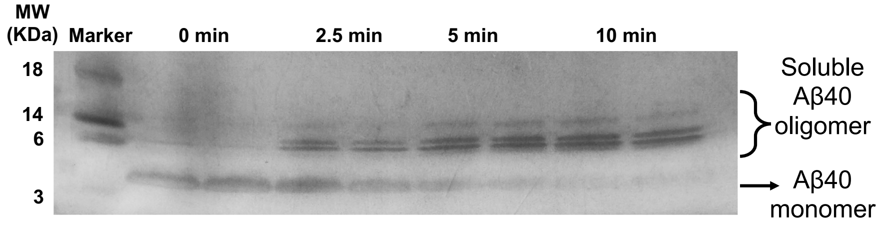


**SI Figure 6.** SDS-Page gel of Aβ40 aggregates with CRANAD-147 under LED irradiation of different durations. The gel was visualized by silver stain.

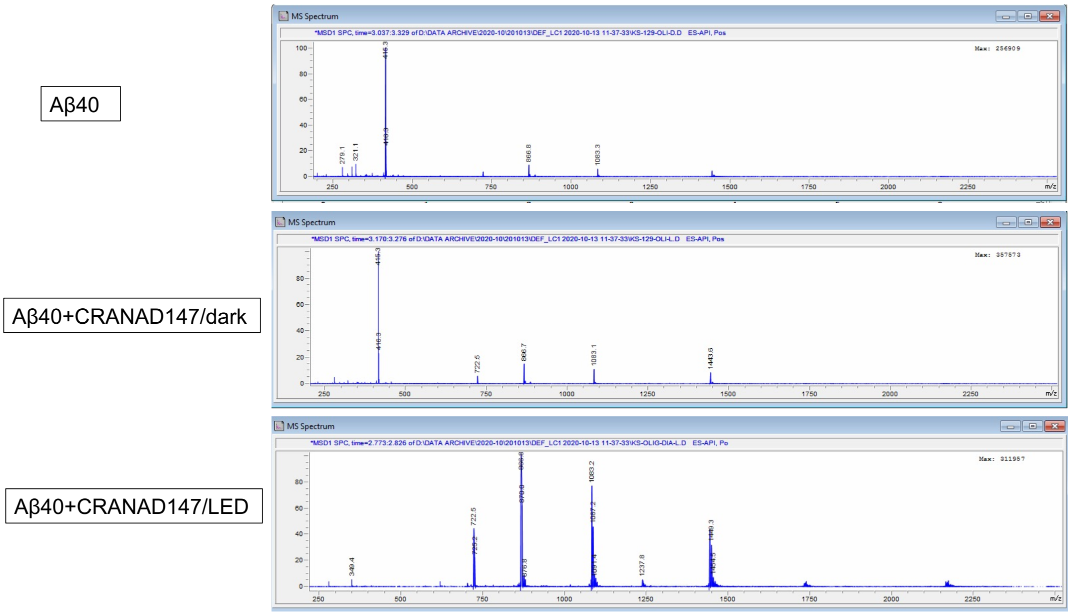


**SI Figure 7.** LC-MS spectra of Aβ40 after different treatments. Several clusters of m/z could be seen in the group of Aβ40/CRANAD-147/LED.

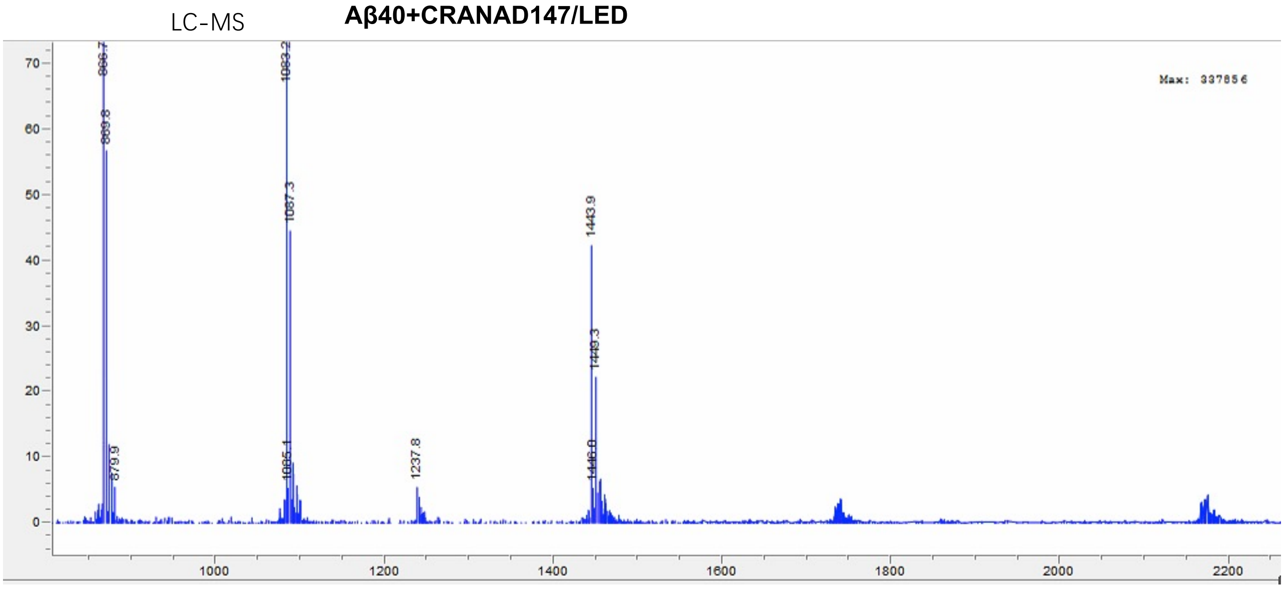


**SI Figure 8.** Zoomed -in MS spectra of Aβ species for the group of Aβ40/CRANAD-147/LED. Several clusters can be clearly seen.

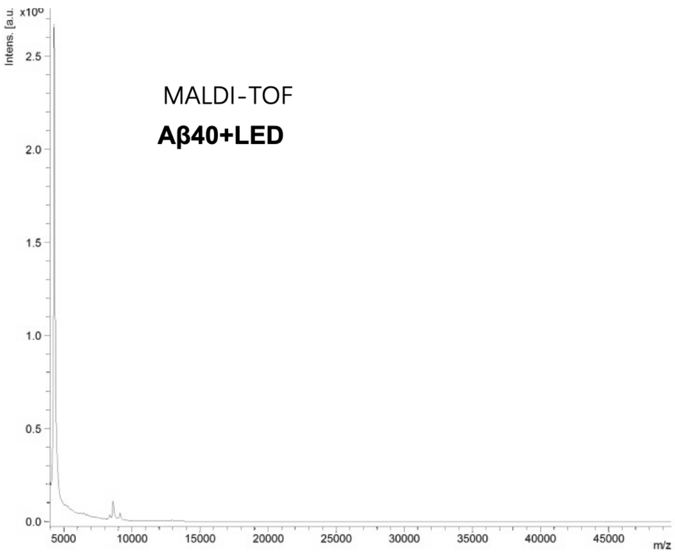


**SI Figure 9.** MALDI-TOF MS of Aβ40 after LED irradiation but without CRANAD-147.

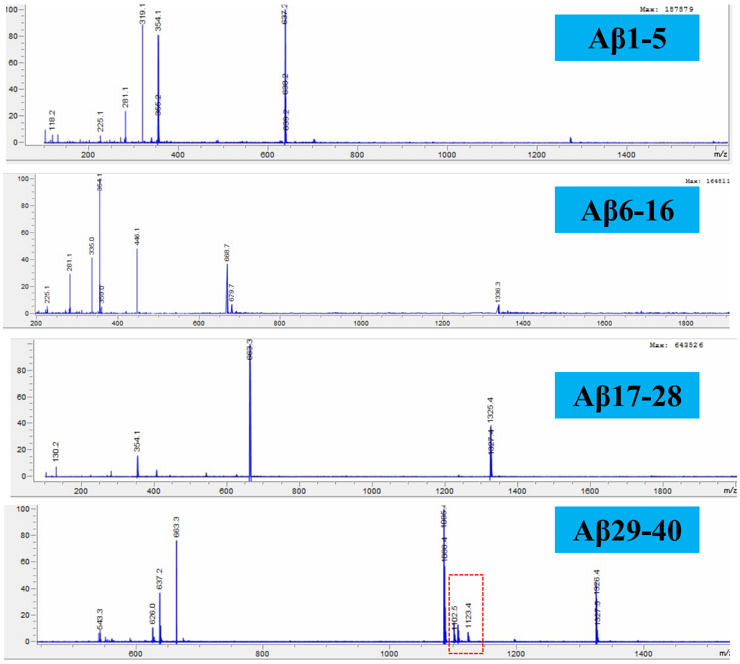


**SI Figure 10.** LC-MS of Aβ fragments from trypsinization treatment of the mixture of Aβ40/CRANAD-147 and LED irradiation.

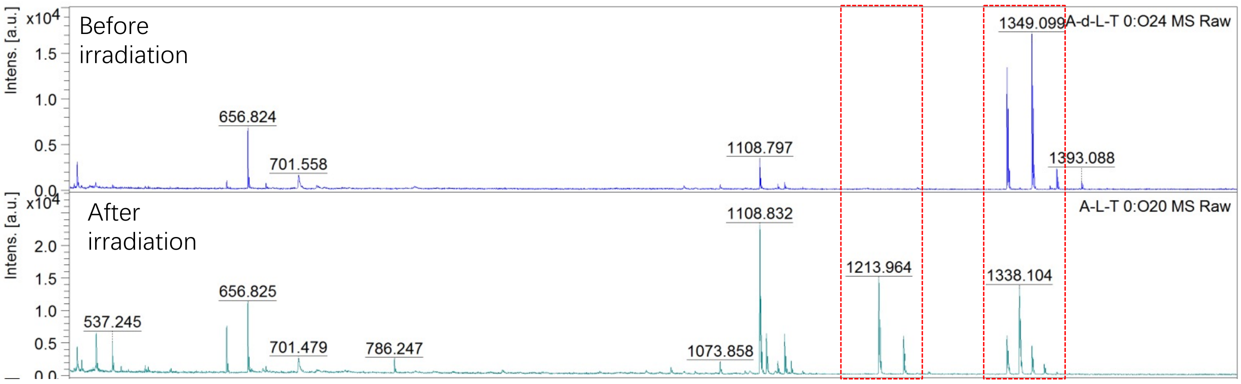


**SI Figure 11.** MALDI-TOF MS of Aβ fragments from trypsinization treatment of the mixture of Aβ40/CRANAD-147 and LED irradiation.

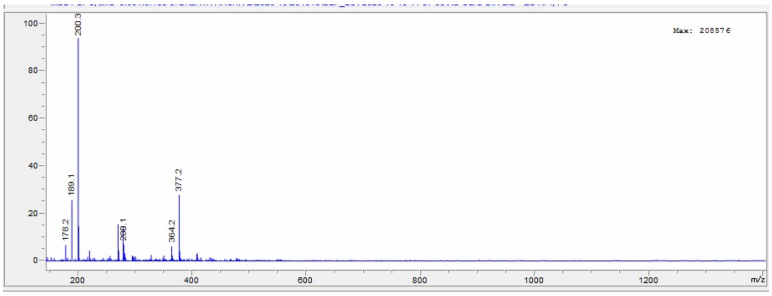


**SI Figure 12.** LC-MS of CRANAD-147 after incubating with ADLumin-4 under a dark environment. m/z= 377.2 is the product of the photo-reaction of CRANAD-147 with molecular light ADLumin-4.

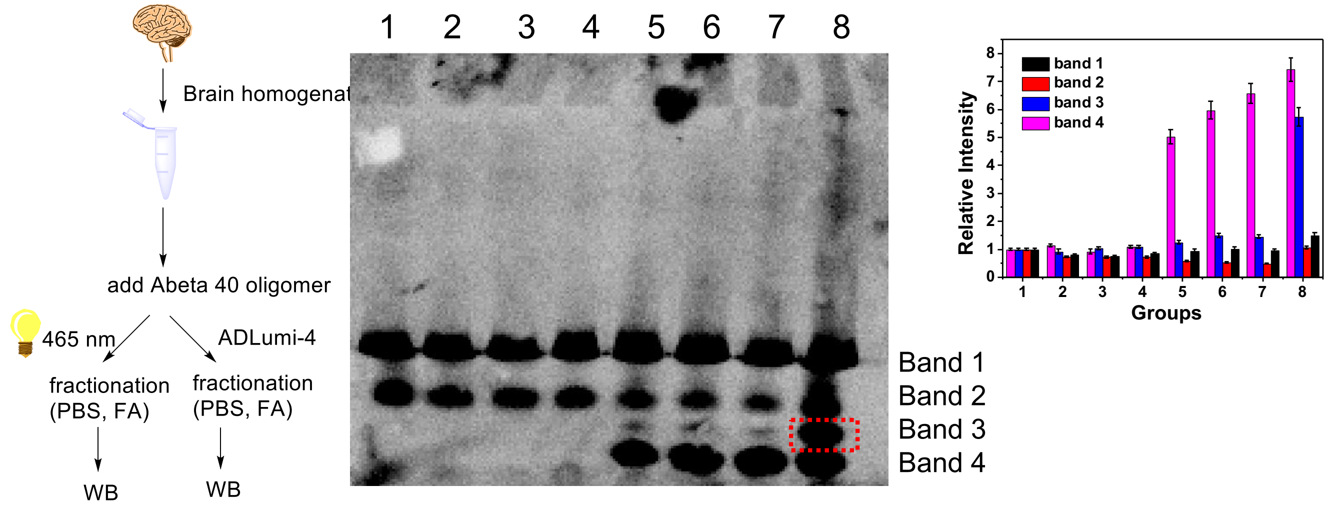


| Group | 1 | 2 | 3 | 4 | 5 | 6 | 7 | 8 |
|---|---|---|---|---|---|---|---|---|
| Aβ 40 | - | - | - | - | + | + | + | + |
| AD-147 | - | - | + | + | - | - | + | + |
| Light | - | + | - | + | - | + | - | + |

**SI Figure 13.** In vitro mimic experiment with externally added Aβ40 aggregates that were treated with CRNAD-147/LED or other conditions. Western blot showed a new band (No. 3) that was absent from groups without Aβ40 addition.

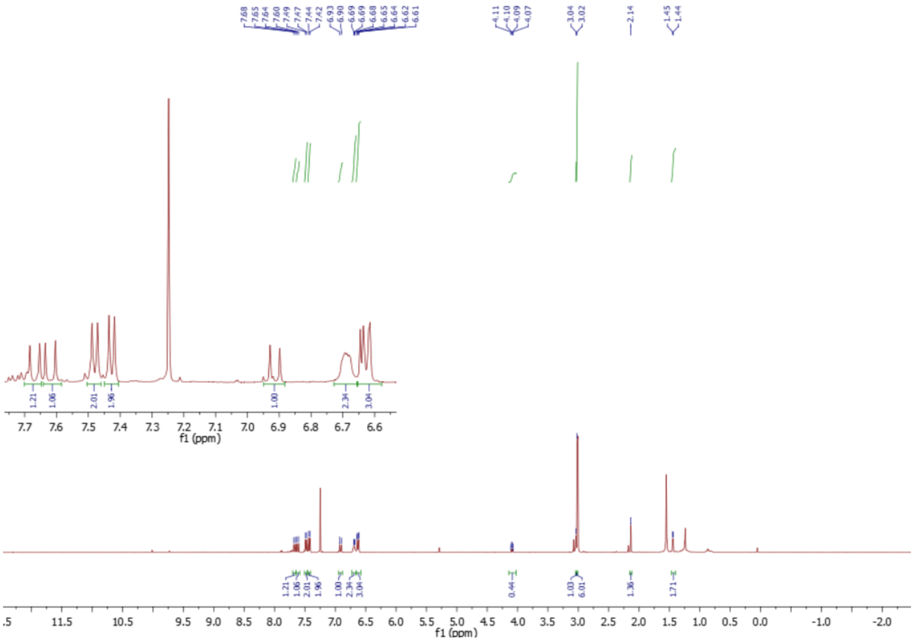


**SI Fig.14** $^1$H NMR spectrum of CRANAD-147 in CDCl3.
